# Mechanical Efficiency Investigation of an ankle-assisted robot for human walking with a backpack-load

**DOI:** 10.1101/2020.11.18.388066

**Authors:** Zhihou Wang, Guowei Huang, Biao Liu, Zikang Zhou, Binghong Liang, Youwei Liu, Longhan Xie

## Abstract

Lower limb assistive robots have a wide range of applications in medical rehabilitation, hiking, and the military. The purpose of this work is to investigate the efficiency of wearable assistive devices under different weight-bearing walking conditions. We designed an experimental platform, with a lightweight ankle-assisted robot weighing 5.2 kg and carried mainly on the back. Eight subjects were tested in three experimental conditions: free walk with load (FWL), power-off with load (POFL), and power-on with load (PONF) for different levels of force at a walking speed of 3.6 km/h. We recorded the metabolic expenditure and kinematics of the subjects under three levels of weight-bearing (equal to 10%, 20%, and 30% of body mass). The critical forces from the fit of the assistive force and metabolic depletion curves were 130 N, 160 N and 215 N at three different load levels. The intrinsic weight of our device increases mechanical work at the ankle as the load weight rises, with 2.08 J, 2.43 J, 2.73 J for one leg during a gait cycle. The ratio of the mechanical work input by the robot to the mechanical work output by the weight of the device decreases (0.904, 0.717, and 0.513 with different load carriages), verifying that the walking assistance efficiency of such devices decreases as the weight rises. In terms of mechanical work in the ankle joint, our results confirm that the efficiency of the ankle-assisted walking robot decreases as weight bearing increases, which provides important guidance for the lightweight design of portable weight-bearing walking robots.

## 1. Introduction

Walking with heavy loads plays an important role in the military (Mala et al.,2015; Gary.,2005) and our daily life (Brian et al.,2016; K E Powell; et al.,1987). Compared with walking without any load, walking with some load has a greater impact on the metabolic expenditure of human body. The index performance is when humans walk at a speed of 1.25 m/s, mass added at the foot has an increasing metabolic cost of 7.5-8.5% per kg (kilogram) during walking, and it is 1-2% for mass added at the waist (Browning et al.,2007). In addition, weight-bearing walking usually causes muscle fatigue, and metabolism will rise as a result (Macdonald et al.,1994; Hogan et al 1998). In fact, the increase in metabolism over a short time will rapidly consume energy of human body. Hence, the risk of injury will rise (Matthew et al.,2011), and the efficiency of work will decrease. Beyond that, long-term weight-bearing walking will lead to damage to lower limb motor function (Grenier et al.,2012).

In recent years, much work has focused on robots assisting human walking with load carriage. The exoskeleton is the earliest and most acceptable type of assist robot. The BLEEX (Berkeley lower extreme exoskeleton) (Zoss et al.,2006) developed by Berkeley University, and factory walking assisting devices (Ikeuchi et al.,2009) developed by Honda, are typical military and civil exoskeletons, respectively. Such devices could reduce muscle fatigue during load-carriage walking indeed. However, the exoskeleton, due to the hard structure itself, would greatly interfere with the users’ natural gait, thus, resulting in great limitations on the research of lowering metabolic cost (Schiele et al., 2009; Schiele et al.,2009; Hidler et al.,2005; Pons et al.,2010). Subsequently, several excellent studies began to focus on assisting single or double lower limb joints with a flexible and wearable device.

Considerable research efforts have been devoted to the improvement of metabolic consumption with the assistance of these wearable devices during walking. An interesting approach in the use of wearable robots for walking assisting was described by CJ Walson et al in 2007. The robot had no motor, only ankle and hip springs and a knee variable-damper (Walson et al.,2007). Although the structure and control method of the robot was modified, the metabolic consumption had not been reduced significantly. In fact, many devices have been shown to be invalid, and may even increase metabolism (Dijk et al.,2011). Nevertheless, these studies have positive guiding significance for subsequent research. For example, during walking, the ankle provides approximately 35% mechanical work, but only accounts for 19% of metabolic consumption (Gorden et al.,2006; Sawicki et al.,2008). In view of such an important effect of the ankle, researchers have begun paying more attention to robot assistance with the ankle, and lightweight design has become a trend (Ferris et al.,2007).

Gradually, some excellent studies have reported a positive decline in metabolic cost. In 2015, MIT designed a no-joint mobile robot that assisted the plantar flexion of the ankle, with its control and transmission mechanism integrated into a small box installed on the shank. The results showed that during subjects’ walking with a 23 kg load, the device could provide an average power of 11.5 W for each ankle joint, resulting in a net metabolic cost reduction of approximately 8±3%. In contrast, the robot can provide an average power of 13.5 W under the condition of no-load walking assist, and the performance was 11 ± 4%. (Mooney et al.,2014; Mooney et al.,2014). In 2014, Alan T.Asbeck at Harvard designed a wearable walking assist robot named ‘exosuit’ that helped humans move the ankle and hip to reduce metabolic expenditure. The designer made use of the transmission characteristics of Bowden cable, instead of using a traditional mechanical transmission structure. The ankle could be provided with an auxiliary force at the peak up to 200 N. It is worth mentioning that the part worn on the leg was much lighter, and this wearable device’s weight distribution implemented the principle of energy efficiency. However, the experimental results showed a 9.3% increase in metabolic expenditure. After that, the auxiliary force increased to 330 N by improving the tension preload. The results showed that, when walking with the load of 30% mass of human body weight, the metabolic cost was reduced by 7.3±5% with the assistance of an exosuit (Asbeck et al.,2015; Paizzolo et al.,2016). Furthermore, in 2017, the Harvard team achieved a 22.83%±3.17% reduction in metabolic cost during walking by placing a very heavy power source and controller outside the body (Quinlivan et al.,2017). Such effect confirmed the great potential of flexible wearable devices for walking assistance. Additionally, their research achieved good results in the field of rehabilitation (Awad et al.,2017).

Previous studies have proven that this type of assisted robot, in the auxiliary process, could replace part of the muscle’s function rather than augment its ability (Meijneke et al.,2014). This gave us an inspiration for our research, whether the autonomous robot from MIT or the exosuit from Harvard, their successful experiment indicates that the performance of reducing metabolic cost was better in assisted load-carriage walking as opposed to no load-carriage conditions, even after taking some other factors into account such as increased tensile length and load. Furthermore, this seems to involve a question of efficiency, since, in any case, the primary goal of achieving a reduction in metabolic consumption is to counteract the effect of the weight of the device on human walking, but the weight of the device is constant, whether weight-bearing or not, so that the same device does the same job, and assistance during weight-bearing can be considered a less efficient situation. It is obvious that the auxiliary force increased, and with no consideration of energy, the performance of metabolic consumption reduction improved as a result (Quinlivan et al.,2017). However, when considering the power of portable devices in the field, the efficiency of the auxiliary becomes very important.

Therefore, it is of great value to explore the efficiency of lower limb assisted robots for evaluating the ability of robot assistance and guiding subsequent research on similar wearable walking assist devices. To achieve this goal, we designed and manufactured a lightweight flexible and wearable ankle-assisted robot. The weight of the device is approximately 5.2 kg, and more than 95% of the weight is placed on the backpack. We used the Bowden cable sheath as the guiding mechanism to transfer the force of the motor to the foot. To obtain experimental data, a gas analysis system was used to record metabolism, a motion capture system was used to record kinematics, and the treadmill was used to maintain a constant gait speed. In our experiments, eight healthy subjects were recorded while walking on a treadmill under three different experimental conditions at a speed of 3.6 km/h.

## 2. Methods

### 2.1. Experimental platform description

When the lower limb moves from the swing phase to the stance phase, the forefoot absorbs energy when the heel contacts the ground to cushion the collision, and store the elastic energy in the Achilles tendon (Kirkendall et al.,2010). In this process, the Achilles tendon in the back foot releases the energy it stores when the front foot hits the ground, helping to shift the body's weight and compensate for the energy loss in the front leg. When the foot leaves the ground, the hip flexors of the hind legs start to drive the lower limbs to swing. Therefore, the elasticity of the tendon is very important to the efficiency of walking, and its releasing energy at the right time can reduce the energy loss due to the heel-strike. During walking, the ankle provides approximately 35% mechanical work, but only accounts for 19% of metabolic consumption (Gorden et al.,2006; Sawicki et al.,2008). From this point of view, assisting the rotation of the ankle seems to be a good option for a walking booster device. For our solution, to avoid affecting the function of energy storage and release of the tendon and to reduce the impact on the gait, we set the timing of the assistance to be between 40% and 60% of the gait cycle, because this is the stage when the ankle does positive work (Collins et al.,2005; McGeer et al.,1990).

The experimental platform, includes the walking assisting robot, a backpack with load, a treadmill, a large frame, a portable gas analysis system (K5b2, Cosmed^®^, Roma, Italy), and motion capture (Vicon^®^, Oxford Metrics, UK; 250 Hz). In the lower actuating part, two kinds of signal acquisition elements are included: a load cell, and a trigger switch. As Fig. 1-b shows, when the robot is on working condition, the driver (Switzerland, Maxon^®^, EPOS4 COMPACT 50/15) receives control instructions that drive the Maxon motor (Maxon EC 647694, 150W, 24V with a 26:1 gear-box) to rotate first, and then the gearbox shaft drives the wire-wheel to rotate to pull the wire rope. Then, the wire rope comes out from the lower end of the Bowden cable sheath, passing through the guide wheel and the limit wheels, in a direction parallel to the bone axis, and the end is connected to the anchor point of the shoe. The load cell (Germany, WIKA^®^, Tecsis F2811) is mounted between this section of the wire rope. At this time, there is an interaction between the robot and human, and the sensor transmitter processed the signal from the load cell and sends it to the signal receiving terminal (German, Beckhoff^®^, terminal bus, EL3124). Then, the real-time tensile force is displayed on the PC screen.

**Fig. 1.**
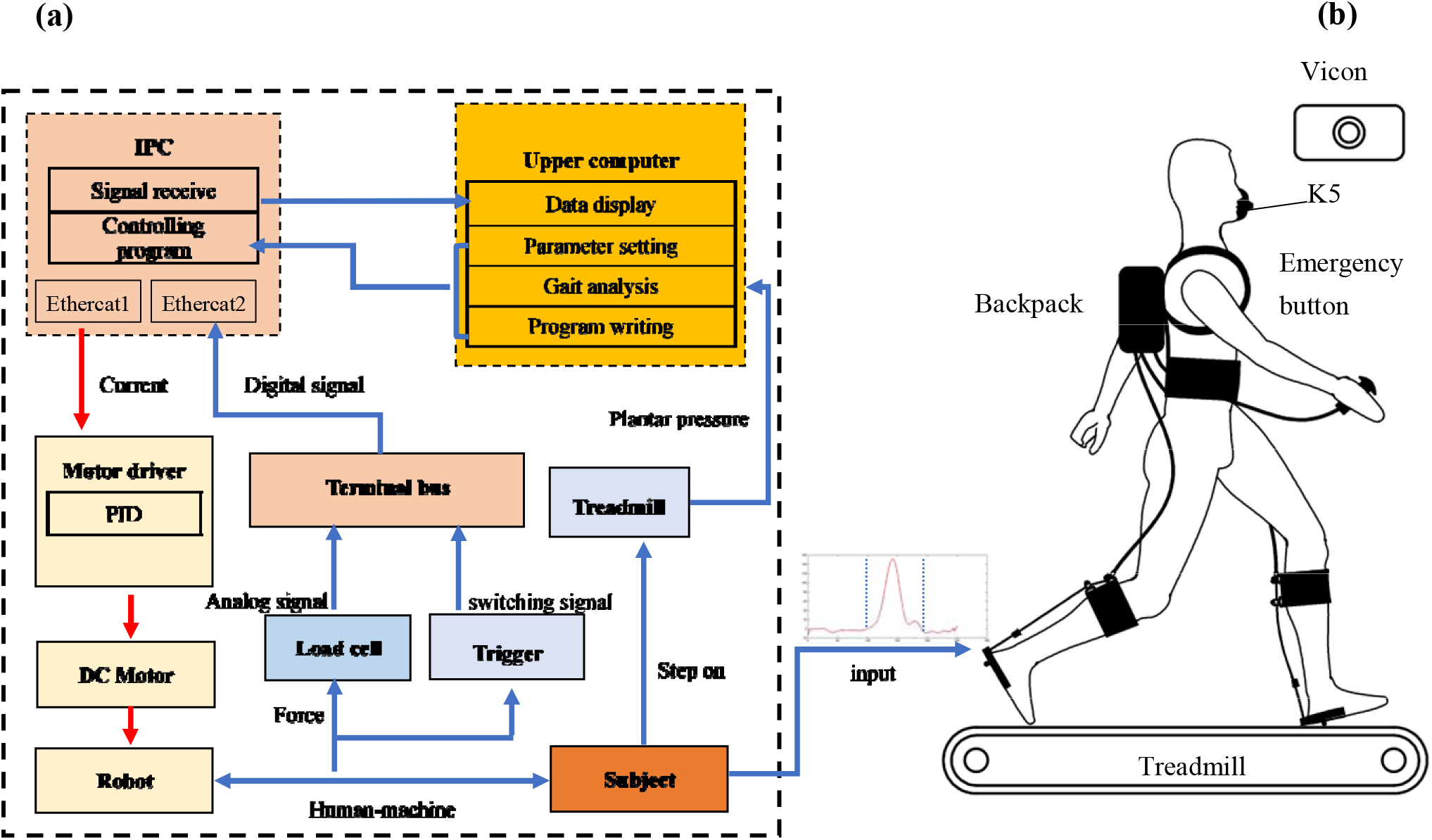
(a) Control system. The complete route of the data stream and power flow of the device. Each block represents a function component of the device, and the function and component of the function component are shown in the block. The red arrow indicates the direction of the power flow from the device, while the blue arrow indicates the direction of the data stream that returns to the device. (b) The overall structure of the experimental platform.

**Fig. 2.**
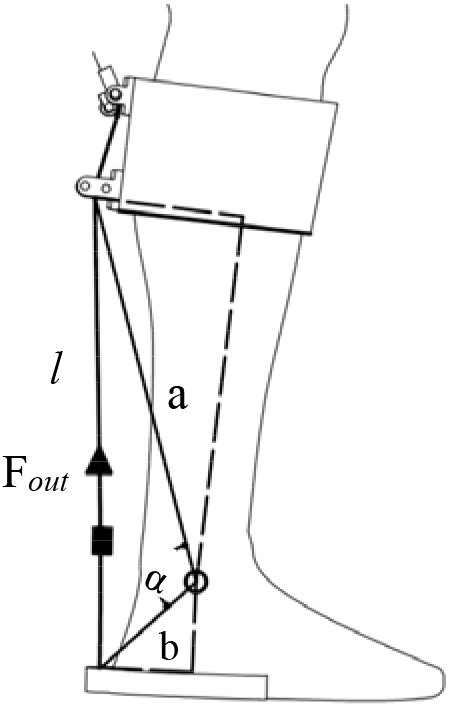
Dimensional models. The circle represents the ankle joint, a, b represents the two lateral sides, *l* is the length of the wire rope, and α is the angle formed by the length of the sides at this time, depending on the moment of gait the body is in.

**Fig. 3.**
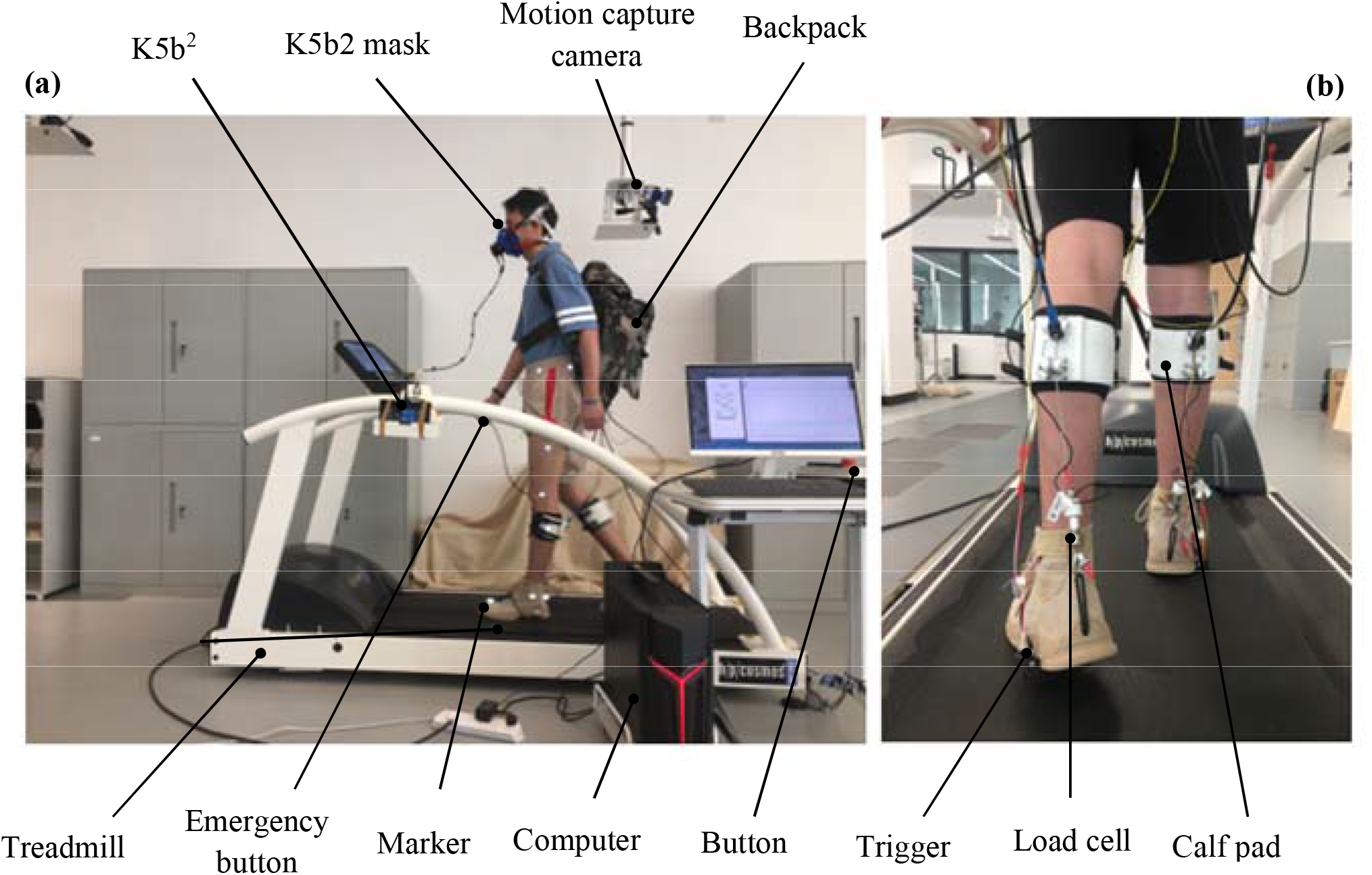
Experimental condition. (a) Side view of the whole experimental plate. (b) Back view of actuator on shank and ankle.

The overall structure of the experimental platform is shown in Fig.1-b. The main body of the device is worn on the human body, the backpack mainly contains the controller and load, and the actuating part is worn on the shank. The treadmill was used to maintain a constant walking speed as the human walked on it. The walking assist robot, mainly includes backpack above the waist and a lower actuator on the calf. The upper actuator, controller (Beckhoff, German, C6015-0010), and load are installed on the backpack together. We set the position of the backpack carried between c1-c12 of the spine, which is conducive to reducing the metabolic consumption of the human body (Stuempfle et al., 2004). The lower actuator, includes the Bowdon cable sheath, lower wire rope, load cell, modified shoes with an anchor, and a soft shin pad (thermoplastic polyurethanes part). As a result, our equipment achieves the goal of minimizing the weight burden on the legs, only adding 0.14 kg of weight (2.7% of the total mass of the robot; the entire robot weighs 5.2 kg) to each leg. When the force is required to be increased, the input current will be added until the target tension is displayed on the PC. During our test, when the subject steps on the treadmill, the trigger switch is pressed, and this time is taken as the beginning of gait. Our equipment primarily assists the ankle in 40-60% of the gait cycle, which is the phase of the gait cycle when the ankle is doing the most positive work.

We use the wire rope as the transmission medium of power, and a flexible sheath of Bowen cable as the guider of transmission. There are many studies on the transmission loss and hysteresis of Bowden cables (Chen et al.,2015; Jeong et al.,2017), The functional relationship between the input and output forces is as follows:

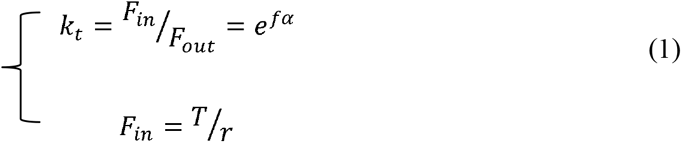

*f* is the friction coefficient, *α* is the wrap angle between two ends, and *r* is the radius of the wire wheel.

According to the characteristics of the DC servo motor, the output torque is:

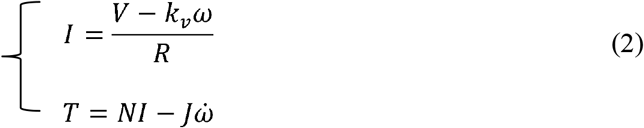

where *I* is the motor current, *V* is the applied voltage, *k*_*v*_ is the voltage constant, *R* is the resistance, *w* and *ẇ* are the velocity and acceleration, *T* is the motor output torque, *N* is the torque constant, and *J* is the motor inertia.

Then the applied mechanical power is estimated by:

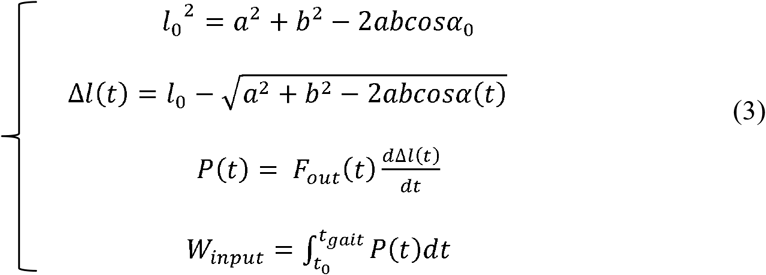

where *l*_0_ Δ*l*_0_, *a, b, α* a represent the length and length change of steel wire, the long and short side of a geometric triangle, and the angle between the two sides. *P* is the power of the robot, *W*_*input*_ is the input mechanical work, and *t*_*gait*_ is the time of a gait cycle.

The change in angle can be converted from the ankle motion captured by the Vicon, and we used the duration of a gait cycle as a criterion for evaluating power. In our calculations, we ignored the wearing part with the deformation of the human body limb, because in approximately 200 N under the action of tension, this deformation is small relative to the change in the length of the wire rope.

### 2.2. Effect of equipment weight on human load-bearing walking

In the introduction, we mentioned that the weight of the device on the metabolic impact, an increase of 1 kg at the waist, metabolism will increase by 1-2%, for the legs, which will be 4-5 times the magnification effect. Flexible wearable devices have the advantage of high efficiency in the structural quality of the hardware. The main mass of the autonomous assistive robot developed by MIT is approximately 2.5kg distributed in the position of the lower legs and feet of the human body (Mooney et al.,2014), which is not an optimal structural design approach from the point of view of efficiency. The Harvard-designed flexible wearable exosuit uses the Bowden line drive to transfer power from the back to the lower limbs of the body. A multi-joint assistive prototype weighs 6.6 kg, with the vast majority of the weight on the back of the body (Asbeck et al.,2015). Therefore, we investigated carrying the weight of the device primarily from the back as a way to improve the efficiency of the assistance.

The quantitative relationships between metabolism and gait, mechanical work, or weight-bearing are inconclusive, and most studies have focused on fitting the mathematical relationships between different variables and metabolism through experimental results (Neputune et al.,2004; Maxwell et al.,2002). In this study, we focus on the variable of mechanical work. The effect of mechanical work of the human body on metabolism satisfies the following mathematical relationship in terms of the mathematical formula. Since the weight of our equipment is not very large in relation to the weight of the load, the calculation from the fitted results has a large error. Because the power from the device is input into the ankle, for output power, we also study the effect of device weight on the mechanical work of the human body from the perspective of ankle-joint power.

Research and technology on human dynamics modeling have been well established (Mcruer et al.,1980), and the inverse kinematic equation can be used to derive the shutdown torque from the kinematic information and plantar pressure of the human body as it walks. Then the power of our joint torque can be obtained by integrating the torque with the angular velocity of the joint motion (Xie et al.,2019). The effect of the weight of the device on the output of human mechanical work during a gait cycle is shown in equation:

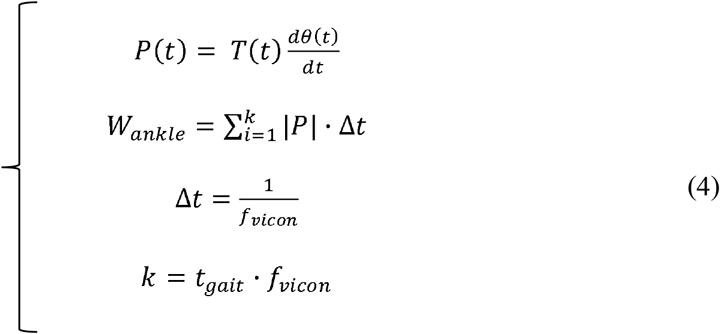

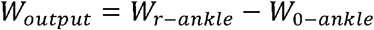

where *P, θ, T,* and *W* represent the ankle moment, angle, mechanical power and mechanical work, respectively, *f*_*vicon*_ is the frame number of motion capture, and *k* is the number of points sampled within a gait cycle.

Previous experimental results have demonstrated that the increase in metabolism is not linear as the load increases, and the rate of increase becomes faster, so it is reasonable to assume that the greater the load class is, the more mechanical work is produced by the weight of the device, at a weight class of 5.2 kg after carrying the load.

### 2.3. Experimental protocol

Eight heathy subjects (22.5±1.5 years; 170.5±2.5 cm; 63.5±3.5 kg) were invited to participate in the experiment. The length of the calf leg circumference, where the TPU parts were worn, the length of the ankle from the foot, and the length of the foot were all recorded for each eligible subject. There were three experimental states in this study as shown in Fig. 5-a: free walking with load (FWL), power-off with load (POFL), and power-on with load (PONL). In the PONL state, one minute before the assist force was applied, the subject need to adjust the walking speed, then, the tension was increased until the target was reached. We divided the size of the auxiliary force into three levels: 150 N, 200 N and 250 N. In each test, we selected one of the three states above, the order of selection was random, and every subject needed to be tested in these three states. After 10 minutes of rest before each test, the subjects were asked to walk on the treadmill at a speed of 3.6 km/h for 6 minutes.

People usually increase walking speed by increasing step length and step frequency. We reminded the subjects to accommodate the speed of the treadmill by keeping the step frequency constant, thus keeping the frequency-dependent metabolic cost constant. We divided the weight-bearing into three grades: 6.4 kg, 12.8 kg, 19.2 kg, about 10%, 20%, and 30% of the body weight. Short-term load-bearing walking would not cause great damage to the joints. Similarly, for the PONL test, we set the range magnitude of the assist force to approximately 50 N. For each set of tests for different loads, the force is approximately 150N at the start level.

Metabolic cost during walking was assessed by K5. Carbon dioxide and oxygen rate were averaged across the last two minutes of each condition used to calculate metabolic power using a modified Brockway equation (Brockway et al., 1987). Metabolic power was normalized by the body mass of each participant. The Vicon was used to record the kinematics, and the treadmill (German, h/p/cosmos, Mercury^®^, FDM-THM,120 Hz) was used to record plantar pressure.

Statistical analysis was conducted in SPSS (SPSS Inc., Statistics 21, USA). One-way repeated-measures analyses of variance (ANOVA) with three modes (FWL, POFL and PONL) were used to verify the effect of the device on metabolic expenditure and mechanical work. Bonferroni post hoc tests were performed to identify differences between conditions when a statistically significant main effect was identified by ANOVA. The significance level was set at p <0.05 for all analyses.

## 3. Result

### 3.1. Mechanical work input

There is no quantitative conclusion as to how mechanical work affects human metabolism, so we use the critical force as our measure of input mechanical work. We assume that there is a force present in a walking-assistive device that exactly counteracts the effect of the weight of the device on human metabolism, where the mechanical work input to the device exactly counteracts the mechanical work output (the effect of the weight of the device), as shown by the fact that the metabolism at this point is equal to the metabolism without the device.

Fig. 4 and Table 1 demonstrate that the weight of the device inevitably increases the metabolic expenditure while walking. Additionally, the greater the auxiliary force provided by the device is, the more pronounced the metabolic decline is, regardless of the load under which the body is walking. Average metabolic power during standing with loads of 6.4 kg, 12.8 kg and 19.2 kg, is equivalent to 10%, 20% and 30% of each participant’s body mass, on average, 1.1±0.1 W/kg, 1.4±0.2 W/kg and 1.6±0.1 W/kg, respectively. The results of net metabolic expenditure under three different experimental conditions are shown in Table 1. The results confirm that, on the one hand, the weight of the device increases metabolic consumption; on the other hand, an increase in the force decreases metabolic consumption. The value of *p* between two sets of experimental data for both effects was less than 0.05.

**Fig. 4.**
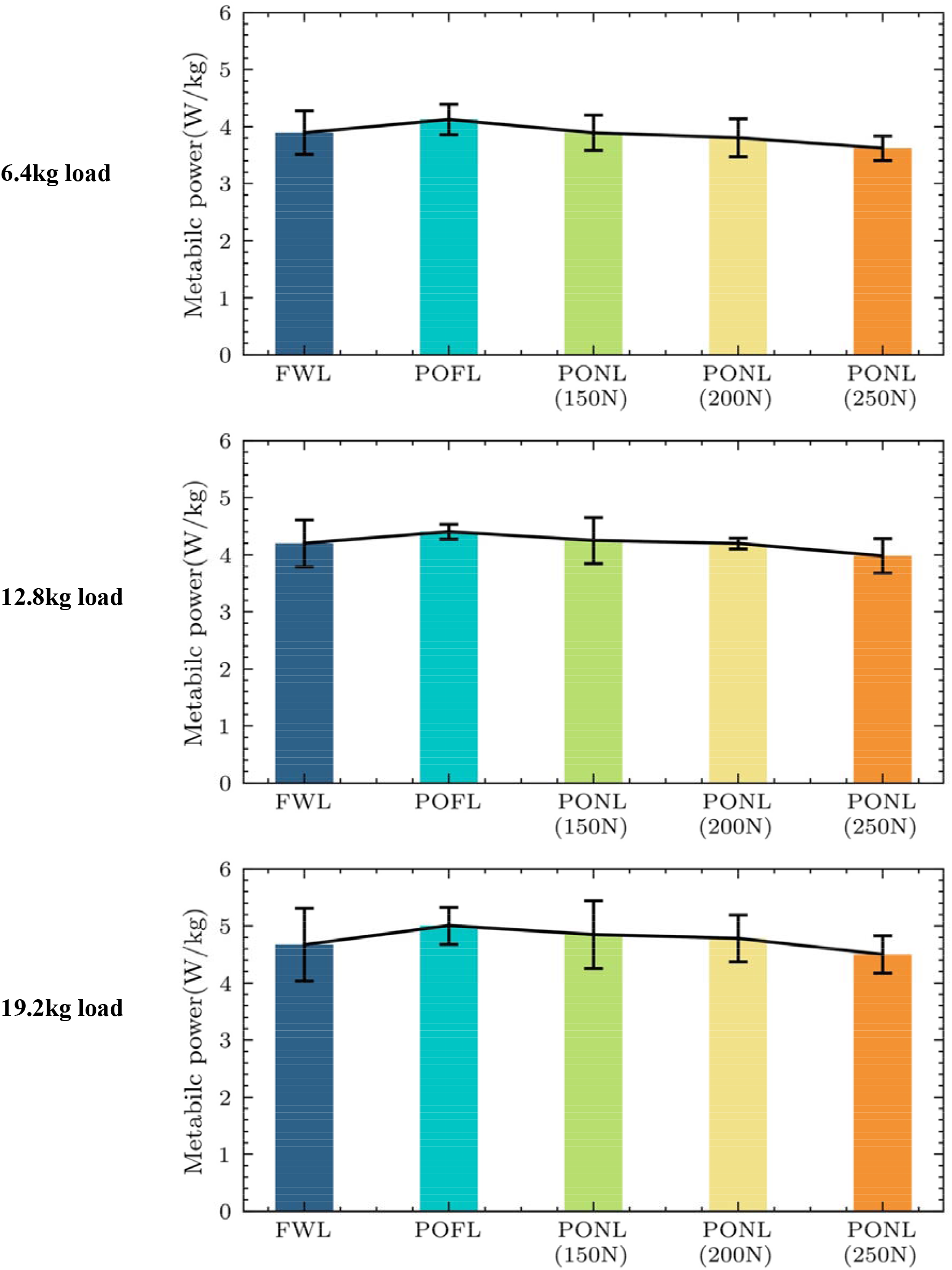
Metabolic power in FWL, POFL and PONL at three different load level: 6.4 kg, 12.8 kg and 19.2 kg.

**Table 1.**
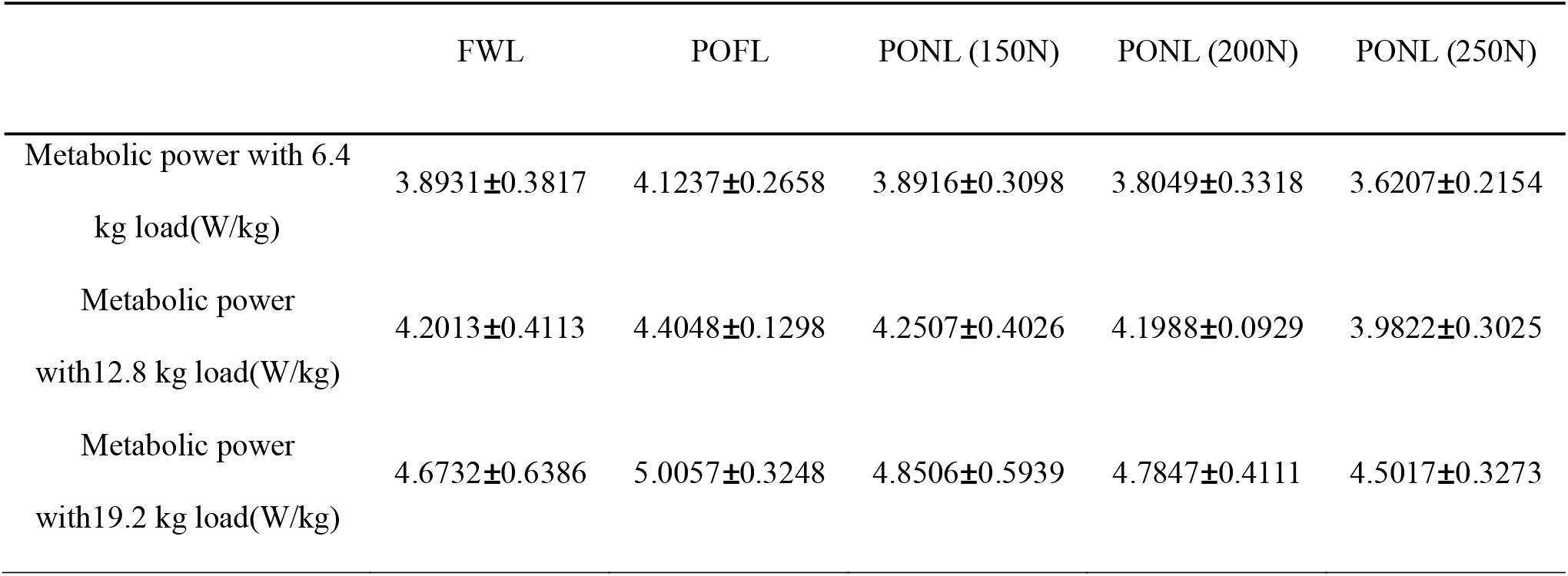
Metabolic power, and gait information for subjects in the experiment

**Table 2.** Mechanical efficiency of assisted robot

| Result \ Load | 6.4 kg | 12.8 kg | 19.2 kg |
| --- | --- | --- | --- |
| Critical force (N) | 130 | 160 | 215 |
| Input work (J/gait cycle) | 2.30 | 3.39 | 5.32 |
| Increased work of ankle (J/gait cycle) | 2.08 | 2.43 | 2.73 |
| Efficiency | 0.904 | 0.717 | 0.513 |

To further obtain the magnitude of the critical force, we performed a linear fit to the force and metabolic cost curves, as shown in Fig. 5. Every goodness of fit is greater than 0.7. The critical force magnitudes are 130 N, 160 N and 215 N for the three different load states. According to the calculation of the input mechanical work in method 2.2, combined with the information about the angle of motion of the human ankle in result 3.1, we obtain the mechanical input work of our assistive device for one leg during a single gait cycle in three load states.

**Fig. 5.**
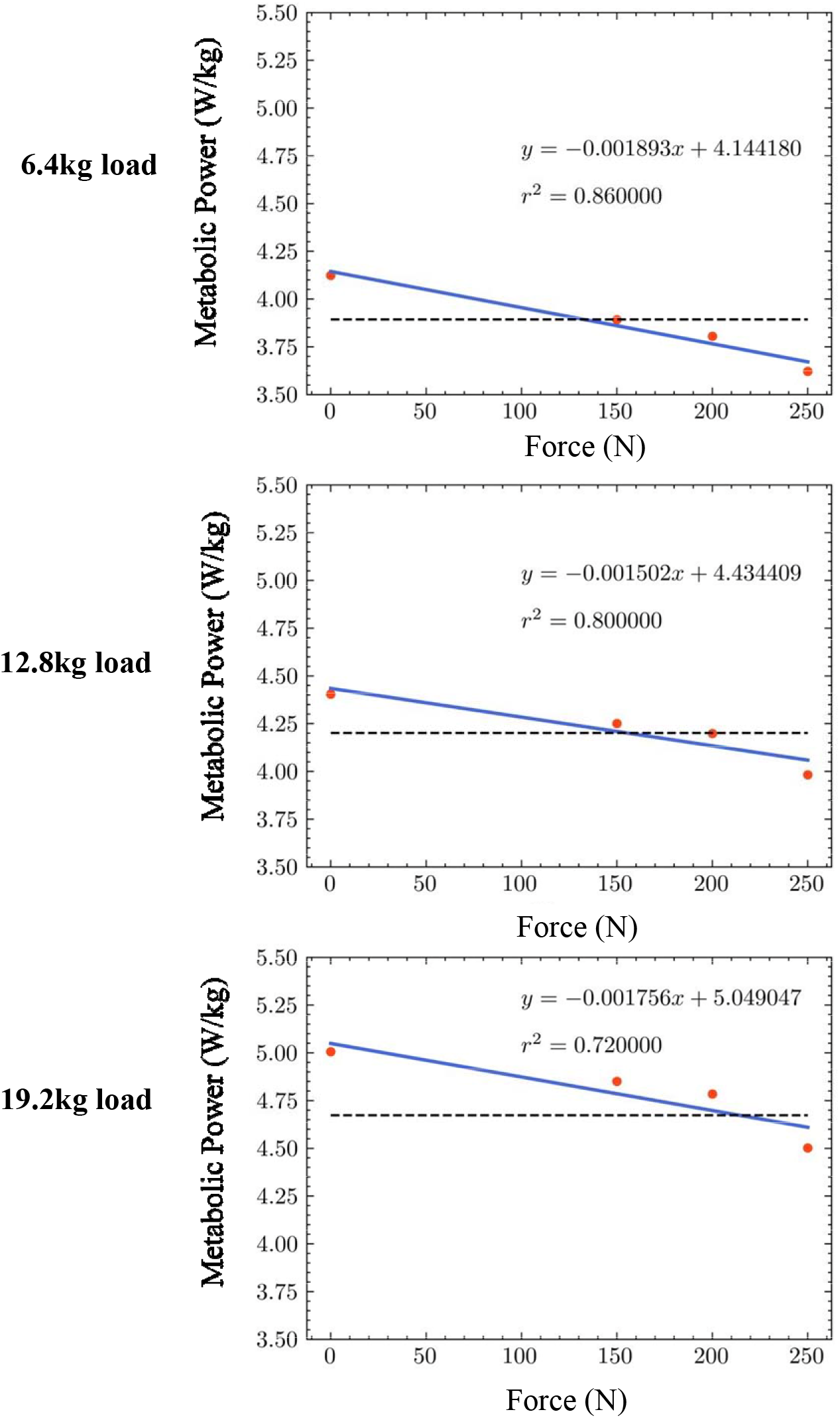
Fitting formula. The red dots represent metabolic cost in this experimental state, the blue line represents the fitted line, and the black dashed line represents metabolic cost in the FWL state. The graphs from top to bottom represent the three load classes.

### 3.2. Mechanical work output

Fig. 6 shows the kinetics of the subject’s ankle in FWL and POFL with different load carriages at 6.4 kg, 12.8 kg and 19.2 kg, and in POFL, a weight of robot at 5.2 kg was added to the human body. The experimental results confirmed that an increase in loading resulted in larger peak angles of ankle-joint toe flexion and dorsal flexion, with 9.1% (1.21 degree) and 18.9% (2.52 degree) increases in the mean peak angle of ankle-joint toe flexion at 12.8 kg and 19.3 kg loading, respectively, compared to 6.4 kg loading. This result confirmed the significance of assisting the ankle for weight-bearing walking. Much energy will be lost during gait phase transition (Graowski et al.2005, Maxwell et al.,2002), and the increase in weight-bearing causes the body to be unable to comfortably provide enough force to push the ankle to perform the gait transition, so it is well understood that energy consumption increases at a faster rate. In addition, the ankle angle data from the POFL group can be used for the calculation of the input mechanical work in Section 3.1.

**Fig. 6.**
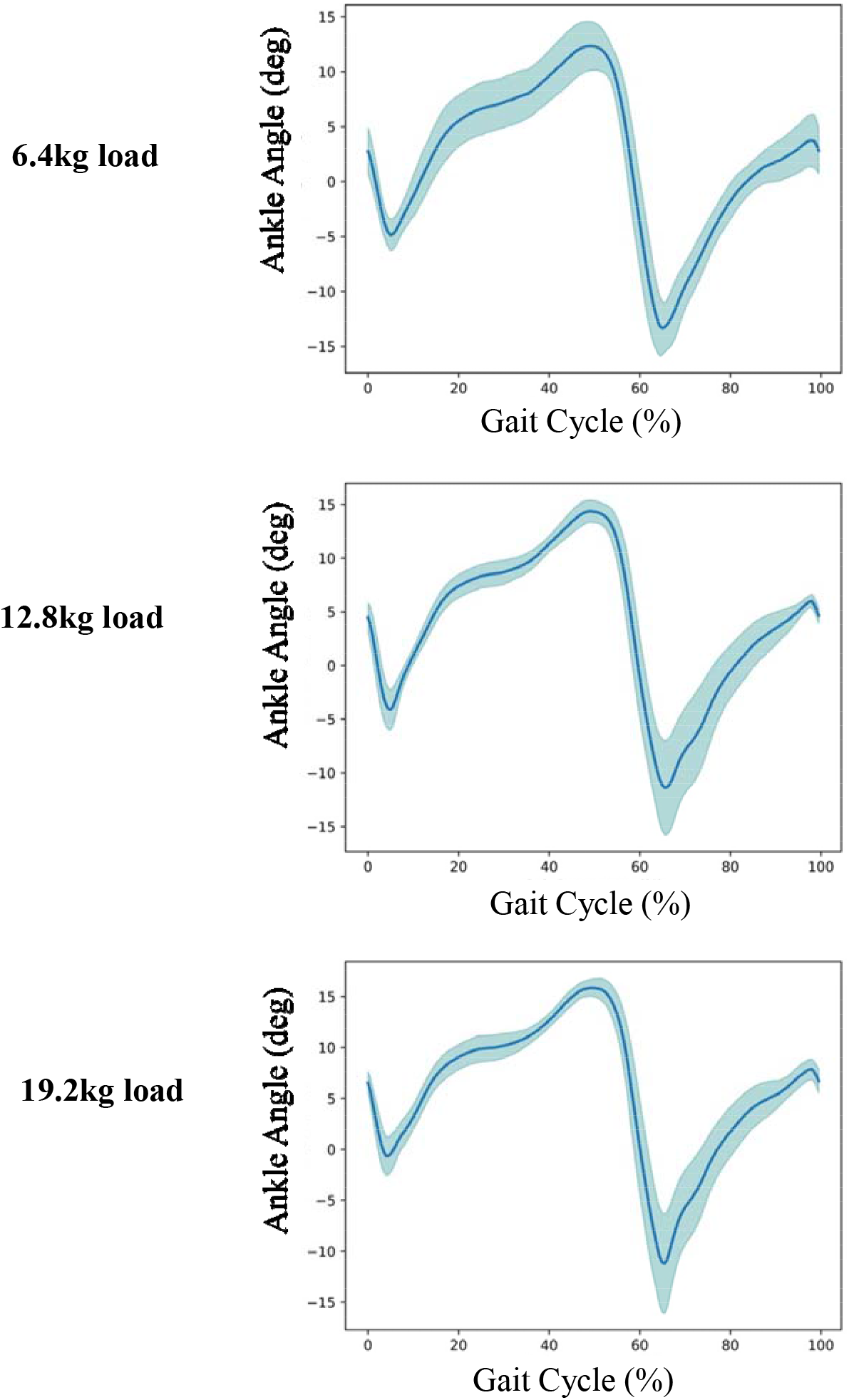
Ankle degree curve in POFL for three different levels of load bearing at 6.4 kg, 12.8 kg and 19.2 kg.

**Fig. 7.**
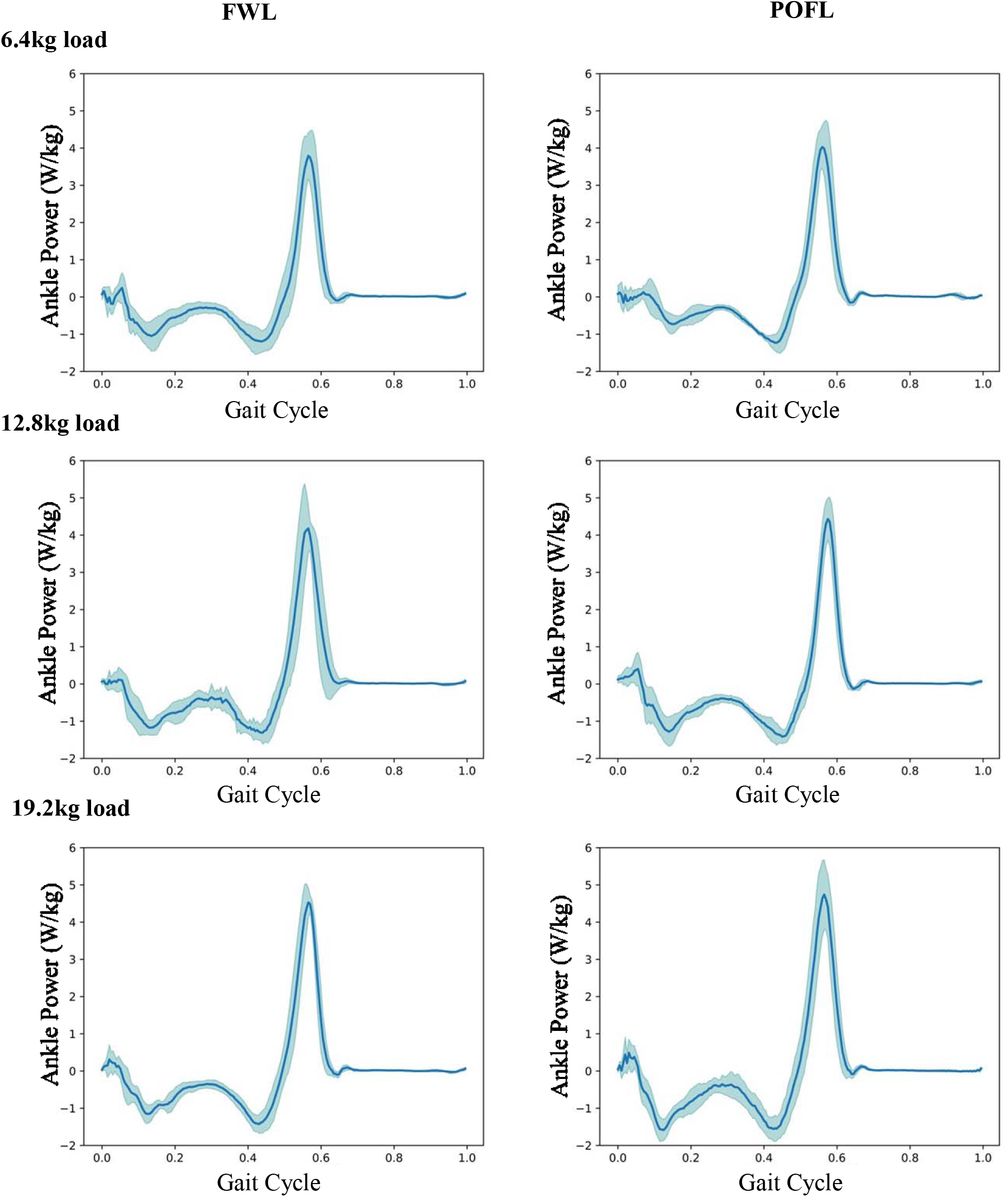
Ankle-power curves in FWL and POFL for three different levels of load bearing at 6.4 kg, 12.8 kg and 19.2 kg. The left side shows the FWL data, and the right side is the PONL data.

From the change in the ankle power curve, it is clear that weight bearing also increases the power output of the ankle joint. The average peak power of ankle flexion at 12.8 kg and 19.3 kg of load increased by 10.23% and 25.02%, respectively, compared to 6.4 kg of load. The robot weight resulted in an increase in the average peak ankle power by 0.2380 W, 0.2521 W, and 0.2134 W at three different load levels. We calculate the magnitude of the ankle-joint mechanical work for the increased weight of the device by using the formula in 2.2.

These test results demonstrate the effect of weight bearing on the kinematics and kinetics of the human lower limb, which is ultimately reflected in the energy metabolic expenditure, and our analysis verifies the significance of assisting the ankle and the robot’s inevitable interference with human walking.

### 3.3. Mechanical efficiency

We start from the ankle joint of one leg to do the work, the input mechanical work is the work done by our assistant robot in the critical force state on the ankle-joint power, the output work is the robotic weight increases the mechanical work of the human ankle joint. The experimental results show that efficiency of the robotic assistance decreases as the level of load increases during the human walking process, and more mechanical work is needed to counteract the adverse effects of the weight of the device on the human body.

## 4. Discussion

This study aims to investigate the efficiency of ankle-assisted robots under different loads during walking. In this research, we designed a robot that could change the load and force level at will, the robot could monitor the real-time tension, and the experimental platform could obtain information about the metabolic cost and human kinematics. From a metabolic point of view, we obtain a critical auxiliary force by fitting a metabolic curve with different assisted forces (where the effect of the auxiliary force exactly counteracts the effect of the weight of the device on the body). Furthermore, the topic of robot mechanical efficiency is raised from the point of view of mechanical work. Experimental results confirm that the mechanical efficiency of robotic assistance to the human body decreases as the load increases.

In fact, as described in the introduction, many studies have investigated the effect of increasing weight and robot assistance on metabolic expenditure. For example, Mooney et al. proposed a formula to calculate the metabolic effect of robotics on walking, through a completely separate perspective from that of assist and load-bearing: 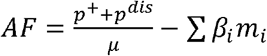 (*p*^+^ and *p*^*dis*^ are the applied positive power and negative power by robot, respectively, *m*_*i*_ is the mass added at different positions of the human body, and *μ* and *β*_*i*_ are weighting coefficients for the booster effect and load impact) (Mooney et al.,2014). In their study, evaluating the effect of the device on human metabolism is also a common way to explore metabolic depletion by fitting the formula through some reviews and experimental results, which cannot truly be quantified (Maxwell et al.,2002). The formula above goes some way to explain the effect of robots on human metabolism, both in terms of assistance and weight gain, which is why we try to avoid robots that add weight to the legs. However, one thing that this empirical formula does not take into account is that the same devices μ and *β*_*i*_ can change when the load is different, taking into account the peculiarities of the human body (Umberger et al.,2011). Therefore, it is not possible to calculate the booster effect and the impact of the load completely separately.

Previous research has suggested that the significance of reducing the weight of a device is to reduce the elevated metabolism resulting from the increased load (Browning et al.,2007), and if this simple linear superimposed relationship can be satisfied, then the same device with the same assistance should have a nearly identical reduction in metabolic consumption under different loads, which clearly is not the case. Therefore, guiding robot lightweight design from a metabolic reduction perspective is a very important point on which focus, but we cannot isolate the link with robotics assistance. Our results confirm that the lightweight design of devices becomes very important in terms of mechanical efficiency. Research on flexible wearable lower limb power-assisted robots presents important guidelines in terms of efficiency and structure. By extension, for mobile portable assistive devices used for hiking or marching, the weight of the device should be of greater concern than for walking assisted robot in wild because of the inconvenience of recharging, which is a great guide to using our limited energy to reduce more metabolic expenditure for the body.

On the other hand, in studying to evaluate the superiority of different devices, our research also provides a new idea. It is naturally good for a device with superior performance to be able to reduce the most metabolic consumption, but one has to consider economic reasons as well. If a device uses less energy to counteract the effects of the device itself on the human body, then the device can be considered superior from an efficiency standpoint.

In addition, in our study, it is worth mentioning that the actuator is mounted on the human calf, and there are counteracting forces that cause discomfort in the calf muscles when the device assists during walking. This flaw needs to be improved later. Moreover, we obtain the critical value of the auxiliary force by fitting, which is logically well understood, but for efficiency, we can only analyze the trend, and not calculate it precisely, because it involves human metabolism, and many problems become very complicated (Umberger et al.,2011).

## Acknowledgements

The authors would like to thank Zhou Zikang for manuscript editing.

## Author contributions

W.Z.H. and H.G.W. provided the conception and design of the research; L.B.H. and L.B. performed experiments; W.Z.H. analyzed data; W.Z.H. and Z.Z.K. interpreted results of experiments; W.Z.H. prepared figures; W.Z.H. drafted the manuscript; W.Z.H., X.L.H., and H.G.W. edited and revised the manuscript; W.Z.H., X.L.H., H.G.W., and L.B. approved final version of the manuscript.

